# Equilibrium morphologies of a macromolecule under soft confinement

**DOI:** 10.1101/2021.05.02.442337

**Authors:** Subhadip Biswas, Buddhapriya Chakrabarti

## Abstract

We study equilibrium shapes and shape transformations of a confined semiflexible chain inside a soft lipid tubule using simulations and continuum theories. The deformed tubular shapes and chain conformations depend on the relative magnitude of their bending moduli. We characterise the collapsed macromolecular shapes by computing statistical quantities that probe the polymer properties at small length scales and report a prolate to toroidal coil transition for stiff chains. Deformed tubular shapes, calculated using elastic theories, agree with simulations. In conjunction with scattering studies, our work may provide a mechanistic understanding of gene encapsulation in soft structures.

---

In recent times, theoretical [1–5], experimental [6–12] and computational [13–19] studies of polymer chains confined in a cylindrical tube has garnered a lot of interest due to a wide range of physical applications [20, 21]. Most studies however, focus on polymer chains under rigid confinement for which conformational statistics are well characterised using scaling theories [1]. In contrast, few studies focus on soft tubular confinement of macromolecules that arise in biophysical problems *e.g.* gene transfer between bacterial pili during conjugation [6, 10], intercellular transport of DNA, [6] and in bioengineering applications such as DNA confined in a lipid tether connecting two giant unilamellar vesicles (GUVs) [8, 9].

Scaling theories of a confined macromolecule in a rigid tube reveal that the axial spread of the chain along the tube axis *R*_∥_ ~ *ND*^−2/3^, where *N* is the number of monomers, and *D* is the diameter of the confining tube [1]. Similar theories have been applied to macromolecules in soft tubular confinement and a phase diagram characterising confined polymer shapes as a function of the surface tension and the size of the chain obtained [4]. Electrophoresis experiments on DNA confined in nanometer sized lipid tubules [9] reveal a “snake” to “globule” conformational transition [4] while Monte Carlo [22, 23] and molecular dynamics [24] simulations have been used to confirm the scaling behavior. These studies however, do not fully capture the effects of semiflexibility and the nonlinear mechanics of the confining tubule on the conformational properties of the macromolecule which sets the stage of the current work.

In this paper we investigate the shape of a deformed lipid tubule encapsulating a semiflexible polymer using molecular dynamics simulations and continuum elastic theories. Our motivation stems from tubular structures observed in biophysical problems. These soft structures have elastic modulii ~ 10 – 20*k*_*B*_*T* such that a confining macromolecule with a size bigger than the box dimension, *R*_*G*_ >> *L*, is able to deform it and results in a shape change.

We consider a bilayer tubular membrane of radius *R*_0_, and length *L* with a constant area *A*_0_ = 2*πR*_0_*L* where *R*_0_ << *L* (SI Fig. 1). The elastic energy of deformation of the tubule is described by the Hamiltonian [25]

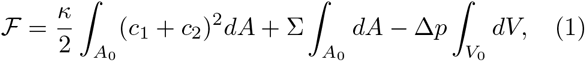

where *κ* and Σ correspond to the bending modulus and the surface tension of the membrane respectively, and Δ*p* denotes the pressure difference acting across the membrane. The energy is expressed in terms of the two principal curvatures *c*_1_, and *c*_2_ (assuming spontaneous curvature *c*_0_ ≈ 0). The undeformed tube radius 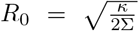 [26, 27] is obtained by setting Δ*p* = 0 in Eq. (1). The deformed tubular shapes are compared against mesoscopic models of bilayer tubules with a confined semiflexible chain.

We perform coarse-grained molecular dynamics (CGMD) simulations of bilayer tubules using the LAMMPS package (SI) following the Cooke-Kremer-Deserno model [29, 40] for a tubule of length *L* = 300*σ*, and different initial radii *R*_0_/*σ* ≈ 9, 11, 13, 15, where *σ* is the unit of length. The bending modulus is obtained by computing the average radius of the tubule ⟨*R*⟩ and the tensile force in the axial direction *κ* = *F*_*z*_*R*_0_/2*π* [29]. This agrees with the value obtained by fitting the radial fluctuation spectrum [30]. The surface tension 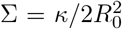, is obtained as a consistency condition from the expression of the tether radius. A semi-flexible polymer is then placed inside the tubule and the coupled polymer-tubule system is simulated over *t* = 5*τ*_*eq*_, where *τ*_*eq*_ = 1×10^4^*τ*_*LJ*_ (where 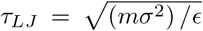, see SI), is the equilibration time. The equilibration time *τ*_*eq*_ is determined by monitoring the radius of gyration *R*_*G*_(*t*) of the polymer undergoing a collapse transition from a swollen to globular state and noting the time at which the *R*_*G*_(*t*) fluctuates about a mean value (SI). The total energy of the system also fluctuates around a mean value beyond *τ*_*eq*_.

Figs. (1) (a) and (b) show the longitudinal and transverse section of a polymer confined in a stiff tubule with *κ* = 24*k*_*B*_*T*. In this regime, *i.e. κ* ≳ 20*k*_*B*_*T*, the entropic pressure due to the polymer cannot deform the tube, and a swollen chain conformation is observed. For softer tubes however, where *κ* ≲ 12*k*_*B*_*T*, the polymer adopts a globular conformation. The conformation of the confined chain as well as the shape of the bulge depends on its persistence length *l*_*p*_. For *l*_*p*_ ≲ *ξ* (Fig. 1 (c) and (d)) the deformed tube has a prolate shape with *R*_∥_ > *R*_⊥_. In contrast for *l*_*p*_ ≳ *ξ*, where *ξ* is spatial deformation scale along the tube axis (Fig. (1)(e) and (f)), the chain adopts a toroidal coil conformation. The bounding tube adopts a non-prolate shape (Fig. (1(f)) with the cross-section akin to a “hockey-puck”. These conformations cannot be captured in the meanfield theories of soft-macromolecular confinement.

**FIG. 1.**
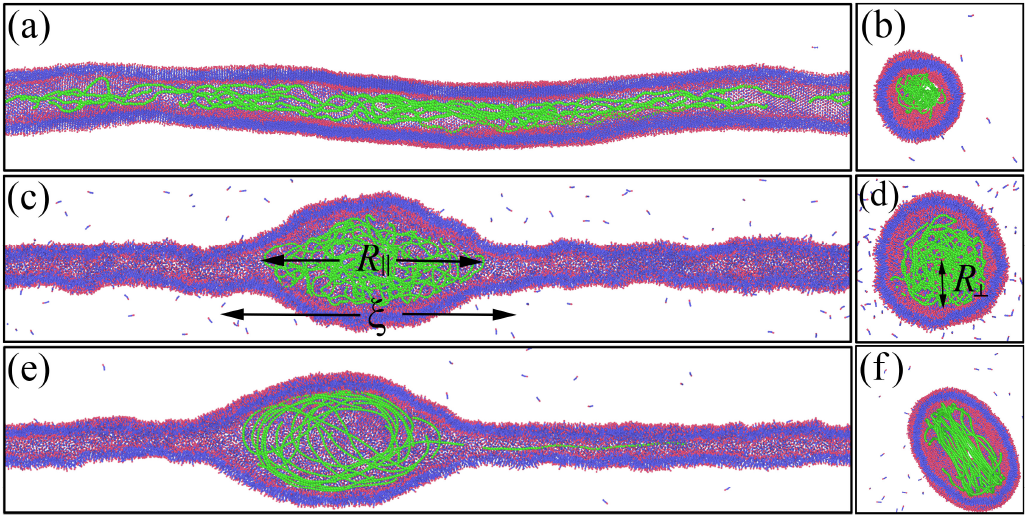
Equilibrium snapshots of axial and radial cross section of a semiflexible polymer (green) of size *N* = 2000*σ*, confined in a bilayer lipid tubule with hydrophilic head (red) and hydrophobic tail (blue) groups obtained from CGMD simulations (SI). Swollen (a) and (b), and globular (c), and (d) chain conformations are observed for rigid *κ* = 24*k*_*B*_*T* and soft *κ* = 12*k*_*B*_*T* tubules respectively. A prolate ellipsoidal (c), and (d), and a toroidal coiled (e) and (f) chain conformation is observed for *l*_*p*_ = 13*σ*, and *l*_*p*_ ≃ 200*σ* such that *l*_*p*_ ≳ *ξ* respectively. (SI Movies.)

We quantify the deformation of the confined tubule on the chain size and stiffness. The time averaged tube radius ⟨*D*(*x*)/2⟩, as a function of the axial distance *x*, with a confined semi-flexible polymer of persistence length *l*_*p*_ ≈ 13*σ* and different polymer lengths *N* = 600, 800, … 2600 is measured, with data collected every 10^3^ steps after equilibration. Fig. (2) shows that the maximum radial bulge ⟨*D*_*m*_/2⟩ of the deformed tubule increases with the chain size, as ⟨*D*_*m*_/2⟩ ~ *N*^0.27±0.02^, for tubules with different initial radii *R*_0_ and 600 ≤ *N* ≤ 7000. The axial deformation length scale *ξ* is independent of the chain length *N* as shown in Fig. (2)(b). Further, the maximum radial deformation ⟨*D*_*m*_/2⟩ is less for tubes with larger initial radii *R*_0_ and a fixed chain size *N*. The deformed tubular shapes resemble a “drop wetting a fibre” [31] (SI), with the axial deformation length scale *ξ* >> ⟨*D*_*m*_/2⟩. It should be noted that the radius of the undeformed segments is less than *R*_0_ enforcing the constant area constraint during the *swollen* to *globule* transition. There is however an associated increase in the volume of the tubule (SI Fig.1 and Fig.7). The effects of semiflexibility on the radial deformation is analysed by computing ⟨*D*_*m*_/2⟩ of a flexible polymer in a tube of initial radius *R*_0_ ≈ 9*σ*. As the radius of gyration of a flexible chain *R*_*G*_ with excluded volume interactions, scales with size as *R*_*G*_ ~ *N*^3/5^*σ* and a soft tubule deforms when *R*_*G*_ ≳ *R*_0_, the lower bound of the size of a flexible chain that deforms the tubule is given by *N* ~ (6*R*_0_)^5/3^. While stiffer chains cause larger radial deformation, we find that it does not alter the scaling exponent.

**FIG. 2.**
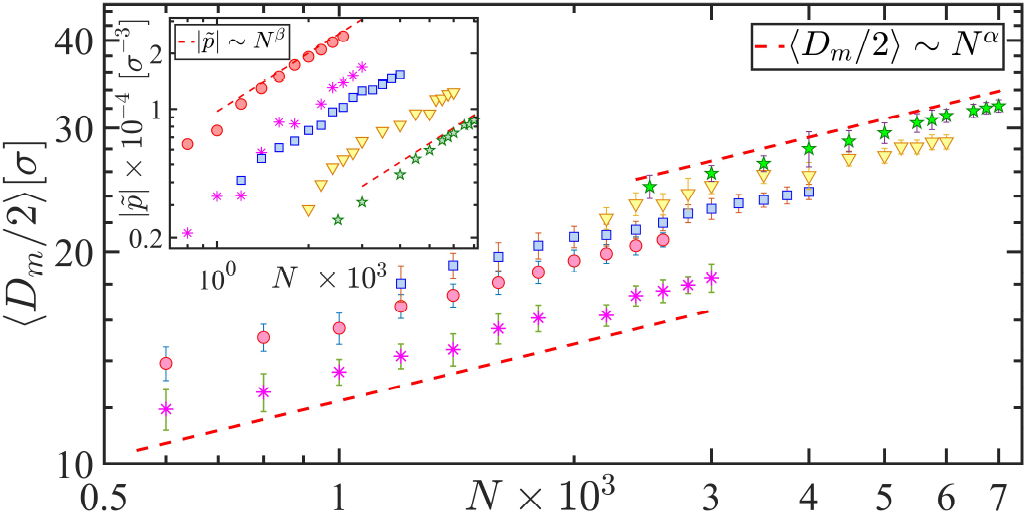
Maximum average radial bulge ⟨*D*_*m*_/2⟩ vs. confined chain length *N* for a semiflexible chain of persistence length *l*_*p*_ ≈ 13*σ* and different undeformed tubule radii *R*_0_ ≈ 9*σ* (⬤), 11*σ* (▇), 13*σ* (▼), and 15*σ* (✶) respectively. A flexible polymer in a confining tube of radius *R*_0_ ≈ 9*σ* (✳), is shown for comparison. Inset shows the variation of the pressure 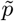 exerted by the polymer on the tubule as a function of *N*, for different radii *R*_0_.

The entropic pressure exerted by the semiflexible polymer Δ*p/κ* scales linearly with the chain length *N* in the strong confinement regime independent of the other parameters, *e.g. R*_0_, *l*_*p*_ *etc.* as shown in the inset of Fig. (2). At equilibrium, the pressure Δ*p/k* is equal to the mechanical pressure exerted by the walls of the confining tubule on the polymer. We use a variational formulation to obtain the pressure exerted by the tubule by parametrising the displacement from the symmetry axis *x* as

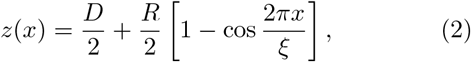

for 0 ≥ *z* ≥ *ξ*. This parametrisation allows us to compute the two principal curvatures *c*_1_ and *c*_2_, as well as the volume and the area of the deformed tubule in terms of *D/R*_0_, *R/R*_0_, and *ξ*. Coarse-graining the deformed tubular profiles obtained from simulations, and using the shape parametrisation we determine the axial deformation scale of the tubule *ξ*. We compute the elastic free energy difference Δ*F* between the deformed tubule and a reference state, a cylinder of radius *R*_0_ and length *L* (*F* = 0) assuming axisymmetric shapes and using the values of *c*_1_ and *c*_2_ in Eq. (1). The pressure exerted by the chain is obtained by equating the free energy difference Δ*F*, to the work done by the chain in deforming the tubule, *i.e.*, Δ*F* = (Δ*p/κ*)Δ*V*, where Δ*V* is the accompanied change in volume of the tubule. For a given value of Δ*p/κ*, we obtain a family of shapes that minimise the free energy. Imposing the constant area constraint we find a unique solution of the shape equations that minimises the elastic free energy. Taken together, this method allows us to obtain the pressure exerted by an encapsulated polymer by solving the shape equation of the tubule (SI).

Polymer conformations inside soft tubules can be characterised on the basis of their spatial extension parallel and perpendicular to the tube axis as a function of the chain length *N*. Fig. (3)(a) shows the variation of the maximum radial extension ⟨*R*_⊥_⟩ for semiflexible polymer with *l*_*p*_ ≈ 13*σ*, as a function of chain length *N* = 600, … 7000. A prolate globular chain conformation is observed and the radial extension scales with the chain length ⟨*R*_⊥_⟩ ~ *N*^0.38±0.03^ in agreement with mean field theories [4]. The inset of Fig. (3)(a) shows the probability distribution of the radial bulge *P* (*R*_⊥_) from which the mean extension ⟨*R*_⊥_⟩ is calculated. The free energy of the confined chain can be computed 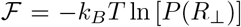. Differentiating the chain free energy with respect to the volume 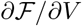 yields the pressure exerted by the chain on the walls. Similar to the confining tubule, the maximum chain extension along the axial direction *R*_∥_ is in-dependent of the chain length *N* as shown in Fig. (3)(b). This is in sharp contrast to the behaviour of confined chains inside rigid tubes where *R*_∥_ increases linearly with chain size.

**FIG. 3.**
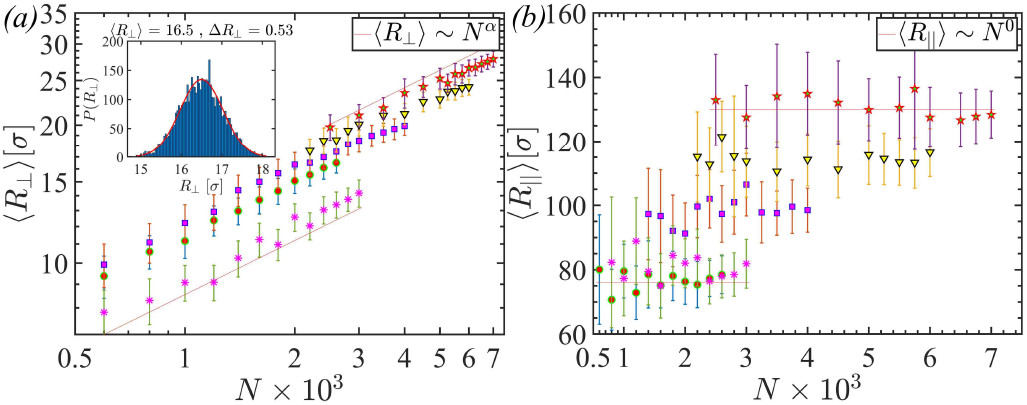
(a) Maximum radial extension of the polymer ⟨*R*_⊥_⟩, vs. chain size *N* for undeformed tube radii *R*_0_ ≈ 9*σ* (⬤), 11*σ* (▇), 13*σ* (▼), and 15*σ* (✶) respectively, compared against a flexible polymer in a confining tube of radius *R*_0_ ≈ 9*σ* (✳). Inset shows the probability distribution *P* (*R*_⊥_) of the radial bulge. (b) Maximum axial extension of the polymer ⟨*R*_∥_⟩ for different undeformed tube radii is independent of the chain size *N*.

While the large scale properties of confined polymer chains can be investigated by computing *R*_⊥_ and *R*_∥_, these do not provide information about confined polymer shapes on length scales smaller than the bulge deformation *ξ*. Fig. (4) shows polymer conformations under various types of confinement. To explore the role of confining geometry, mechanical stiffness and solvent quality on the shape of a confined polymer we compute the *(a)* tangent-tangent correlation function 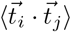, *(b)* asphericity parameters Δ, and Ξ, *(c)* mass distribution *ρ*_*r*_ as shown in Fig. (4). Space filling pictures of four compact conformations, (a) spherical, (e) prolate, (i) toroidal coil and (m) collapsed prolate shapes are shown. These statistical quantities that explore the short distance properties of the confined chain are used to characterize different compact shapes.

**FIG. 4.**
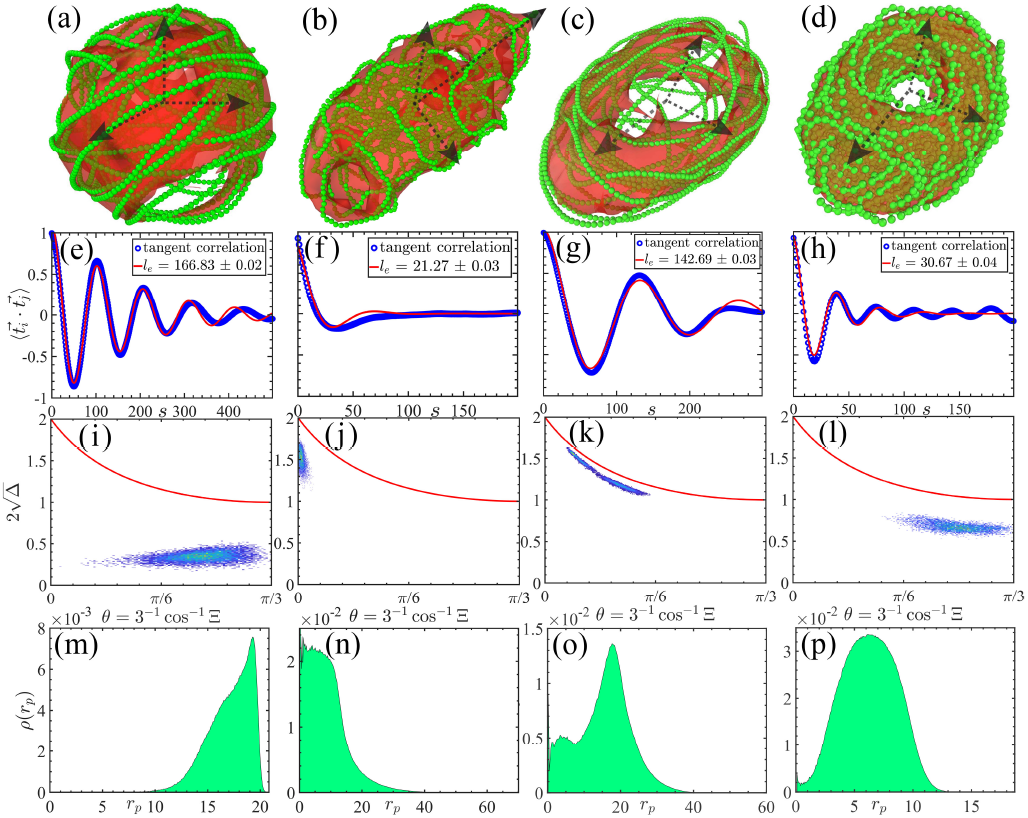
Conformations of semiflexible polymers: (a) inside a rigid sphere with *l*_*p*_ ≈ 200*σ*, a soft tube with *κ* ≈ 12*k*_*B*_*T* and (b) *l*_*p*_ ≈ 13*σ*, (c) *l*_*p*_ ≈ 200*σ*, and (d) *l*_*p*_ ≈ 200*σ* in a bad solvent. Panels (e)-(h) shows tangent vector correlation function of the polymer obtained from simulations (blue) and fit to a functional form (red) (see text). For stiff chains *l*_*p*_ ≈ 200*σ*, a shape transition from ellipsoidal to toroidal conformation is observed, characterised by the asphericity parameters, Δ, and Σ. Distribution of monomers from the centre of mass of the chain (panels (m)-(p)), are used to distinguish chain conformations baed on the confining geometry, polymer stiffness, and solvent quality.

The tangent-tangent correlations of a semiflexible polymer under hard spherical confinement is given by 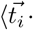 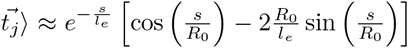 up to 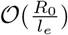 [32]. The leading order contribution to the correlation function for 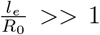 is given by 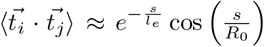, which we have used to fit the tangent vector correlation data obtained from simulations. In our simulations the radii of the confining sphere and the undeformed tubule are *R*_0_ ≈ 20*σ*, and *R*_0_ ≈ 9*σ* respectively. Fig. (4)(e)-(h) shows the fitted form against simulation data. The effective persistence length *l*_*e*_ obtained from the fit is is smaller than the bare value in free space for stiff chains. For example, Fig. 4(e) shows the tangent-vector correlation function fitted against simulation data which yields an effective persistence length *l*_*e*_ ~ 167*σ*. This can be explained simply by noting that inside the rigid spherical cavity, a stiff-polymer with *l*_*p*_ ~ 200*σ* >> *R*_0_ wraps around the walls. As a result, the tangent vectors are decorrelated over *πR* ≈ 62*σ*, *i.e.* opposite points of the semicircular arc traced by the polymer. The effective persistence length *l*_*e*_ for semiflexible chains in contrast is greater than bare persistence length *l*_*p*_ ≈ 13*σ* under soft confinement (Fig. (4)(f) *l*_*e*_ ≈ 20*σ*) and in a bad solvent (Fig. (4)(h) *l*_*e*_ ≈ 31*σ*). The increase in the persistence length is due to the limitation of the functional form used to fit the correlation function. The periodicity in tangent vector correlations (Fig. (4)(e)) is due to the rigid constraint enforced by the wall. A similar oscillatory behavior is observed for soft confinement of stiff polymers *l*_*p*_ >> *R*_0_ (Fig. 4(g)) with the polymer forming a toroidal coil. A collapsed conformation of the confined polymer in a bad solvent (Fig. 4(h)) with *l*_*p*_ ≈ 13*σ* also shows an oscillatory behavior. This feature is absent under soft confinement (Fig. 4(f)) where *l*_*p*_ ⪆ *R*_0_.

The shape transition can be quantified further using the asphericity parameters 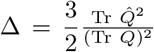 and 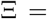 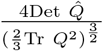, expressed in terms of the variance of the eigen-values of the traceless gyration tensor about the centre of mass, 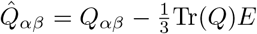, where *E* is a unit tensor [33]. The gyration tensor, 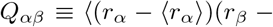 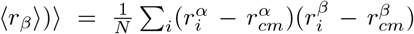, where *α* and *β* indicate components of the position vector of the *i*-th monomer [34, 35]. The average chain length is represented by 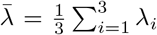. The asphericity parameter Δ lies in the range 0 ≤ Δ ≤ 1, with Δ = 0 for spheres, and Δ = 1 for rigid rods. The difference between oblate and prolate shape is captured by the parameter Ξ. As shown in Fig. 4 panels (i)-(l), the heat map obtained for different configurations in the Δ-Ξ phase plane [34] is distinct for different shapes. The red line in the figures demarcate the region between accessible and inaccessible closed shapes, while the heat map provides a measure of prolate, and oblate polymer conformations, which differentiate between ellipsoidal and toroidal shapes Fig. (4)(j) and (k) respectively.

Further, we compute the monomer distribution 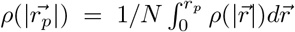, as a function of *r*_*p*_, where 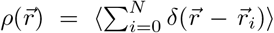, is the local density. Fig. 4 (m) shows 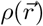 for a rigid spherical confinement of radius *R*_0_ ≈ 20*σ* has a maximum at *r*_*p*_ ≈ *R*_0_ corresponding to the stiff chain wrapped inside a spherical cavity. In contrast, for soft tubular confinement (Fig. 4(n)), the monomer distribution has a peak at *r*_*p*_ ≈ 0 for globular chain conformations with stiffness *l*_*p*_ ≈ *R*_0_. For stiffer chains *l*_*p*_ >> *R*_0_ where the chain conformation is a toroidal coil exhibits a bimodal distribution with peaks at *r*_*p*_ ≈ ⟨*D*_*m*_/2⟩ and *r*_*p*_ ≈ 5*σ* corresponding to the thickness of the toroidal shell estimated from volume conservation (see Fig. 4 (o)). A similar shift in the peak of the monomer distribution arises for collapsed chains in a bad solvent due to attractive interactions. The monomer distribution is however broader (Fig. 4(p)) in comparison to the toroidal coil. These quantitative statistical measures provide a benchmark by which semiflexible polymer conformations under soft-confinement can be characterised.

In summary, we have developed a multiscale model to characterise conformations of polymer chains under soft confinement. These confined phases are a function of the polymer chain length *N*, its persistence length *l*_*p*_, the radius of the undeformed tube *R*_0_, and its bending modulus *κ*. For soft tubules *κ* ≲ 12*k*_*B*_*T* we observe a swollen to globular conformational transition as the chain size *N* and stiffness *l*_*p*_ is increased. As opposed to globular structures reported in previous scaling studies [4] we find that a polymer adopts ellipsoidal conformations with radial bulge of the polymer ⟨*R*_⊥_⟩ increases with the chain size. The maximum radial bulge of the deformed bilayer tubule follows a different power law scaling with the chain size *N*.

It is difficult to resolve macromolecular conformations inside soft tubules of nanometer dimensions using conventional scattering methods in experiments. In such situations our calculation scheme can be used to parametrise the deformed tubular shape and obtain the principal curvatures and the change in volume (Eq. (1)). Further, a measurement of the bending modulus of the tubule *κ* from flicker spectroscopy allows us to compute the energy of the deformed tubule. This allows us to compute mechanical pressure exerted by the polymer on the walls and in turn obtain the conformational statistics of a deformed chain in terms of the radial and axial chain deformations.

In addition, we have identified statistical measures characterising deformed polymer conformations on length scales smaller than the deformation scale in terms of the aspherity parameters, the monomer distribution and the renormalised persistence lengths. These measures can be used to distinguish deformed tubular conformations that have similar shapes on length scales comparable to the axial deformation *ξ*. As an illustration, we report a novel ellipsoid to toroidal chain conformational transition as a function of the persistence length *l*_*p*_ inside soft tubes. We hope that our theoretical work will motivate experimental studies in this direction which will in turn shed light on biophysical problems involving soft macromolecular confinement.

## ACKNOWLEDGMENTS

SB and BC thank University of Sheffield, IMAGINE: Imaging Life grant for financial support. We thank Dr. B. Mukherjee, for a critical reading of our manuscript and Dr. S. Kundu for developmental edits.

## I. APPENDIX

## II. SHAPE EQUATION OF TUBULES

## A. Tubular lipid membrane

We develop a variational formulation to predict axisymmetric deformed tubule shapes (altered via membrane insertion) by minimising the membrane free energy [25, 36, 37]. A constant pressure Δ*p* acts across the membrane. Motivated by our simulation results, we work in an ensemble with the constant surface tension Σ and pressure difference Δ*p*. The elastic energy to deform a tubular membrane accounting for changes in area and volume, is given by the Helfrich-Canham free energy [25]

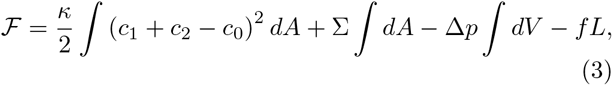

where *c*_1_ and *c*_2_ are the the two principal curvatures of the axisymmetric tubule of length *L* and *c*_0_ is the spontaneous curvature set to zero in our simulation. The energetic cost of pulling out a tether of length *L*, from a GUV, relevant for micropipette aspiration experiments is modelled by a tensile force *f*. For a cylindrical tubule, the principal curvatures 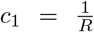, and *c*_2_ = 0. The elastic free energy of the tubule is given by, 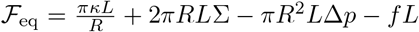. The equilibrium shape corresponds to a zero pressure difference acting across the membrane, *i*.e., Δ*p* = 0. The shape equations result from a variational minimisation of the free energy, *δF* = 0 and leads to the shape equations 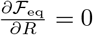 and 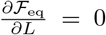. The equilibrium radius of the tubule, 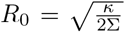 and tensile force acting along the tube axis 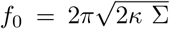 can be obtained from the shape equations [26].

Solutions to variational shape equations for axisymmetric tubules with open boundaries for tubular geometries have been explored, though differing opinions regarding the appropriate boundary conditions have been reported [38, 39]. Fig. (5) shows a schematic figure of an axisymmetric bilayer tubule deformed by an encapsulated polymer chain. We define a parameter 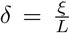, where *ξ* is the axial deformation of the tubule. The extent of deformation can be quantified by comparing the dimensionless ratio *δ/ρ* where *ρ* is the aspect ratio of the undeformed tubule *ρ* = *R*_0_/*L*. Minimization of the curvature energy Eq. (3) with appropriate boundary conditions incorporating geometric and physical constraints <u>e.g.</u> constant area, volume, *etc.*, imposed by the physical situation yields the equilibrium shape of deformed axisymmetric tubules.

**FIG. 5.**
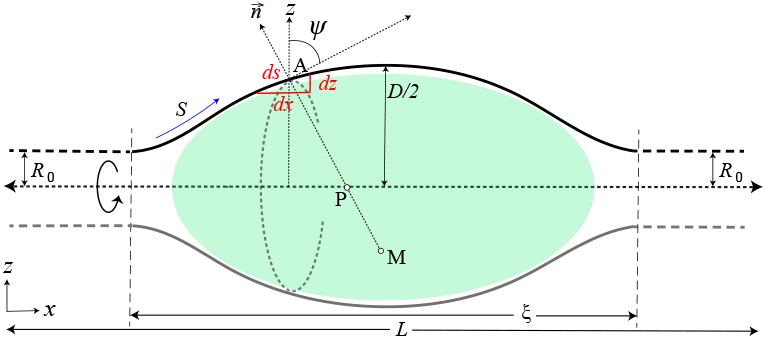
Schematic representation of the cross section of a deformed tubule in the *X* − *Z* plane with the tube along the *X* direction. An axisymmetric shape is assumed. Tube deforms in the plane perpendicular to the *X* axis. The angle between the tangent at a point (e.g., A) on the curve and the *Z* axis is denoted by *ψ*(*S*), where *S* is the arc-length of the tubular contour.

Fig. (5) shows the principal curvatures 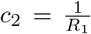 and 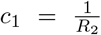 of a deformed tubule at point *A*. At this point, *R*_1_ = AM corresponds to the radius of curvature along the meridian curve (M being the centre of the meridian curve in the plane of the figure). The radius of curvature *R*_2_ = AP along a direction perpendicular to the tube axis is centred at point P which lies on the axis of rotational symmetry along *X* direction. From the parametrisation of the surface we obtain *dS* = −*R*_1_*dΨ* and *Z* = *R*_2_ sin *Ψ*, and the geometric relations 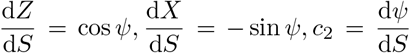, and 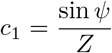 (Fig. (6)). We express the elastic free energy of the membrane tube in Eq. (3) using the above parameterisation, 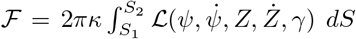, where the <u>Lagrangian</u> is given by [37]

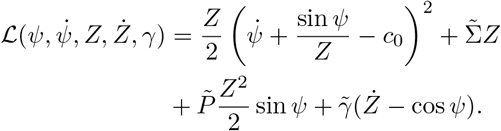

**FIG. 6.**
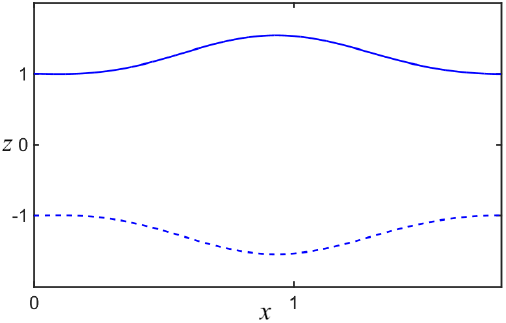
Cross-section of a deformed axisymmetric tubular membrane obtained by solving Eqs. (4a) - (4c) with the initial conditions 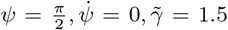, *Z* = 1, 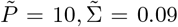. We start the integration at *S* = 0 (as there are no divergent term for *Z* ≠ 0) and stop the integration at *S* = 3 where *Ψ* reaches at 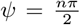. The dotted line is the reflection of the solution about the mid-plane of the tubule.

The first variation of the free energy in Eq. (3) yields one second-order (expressed in terms of two nested first-order differential equations) and two first-order differential equations in terms of the variables Ψ(*S*), *U* (*S*), 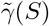, and *Z*(*s*) [37] given by

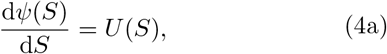

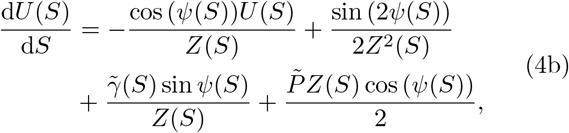

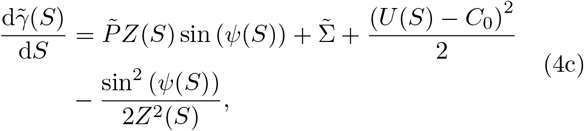

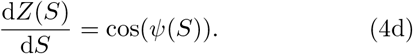

We solve the coupled differential equations Eq. (4) with initial conditions 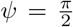, and 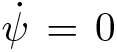. In our numerical scheme we integrate Eq. (4) starting from *S* = 0 corresponding to Ψ(*s*) = 0, till *S* ~ 1, at which Ψ = *π*/2. This corresponds to a point *X* = *X*_*m*_ along the tube axis. The profile is then reflected about a line perpendicular to the *X*-axis to generate the membrane configuration along the arclength *Z*(*s*) above the midplane of the tube axes as shown in Fig. (6). A dotted line showing the reflection of the membrane profile obtained by reflecting *Z*(*S*) about the midplane of the tubule is also shown. Taken together, Fig. (6) represents the cross-section of the axisymmetric deformed tubule. We have compared equilibrium axisymmetric tubular shapes obtained from a minimisation of the curvature energy with those obtained from coarse-grained molecular dynamics simulations for specified value of surface tension Σ and pressure difference Δ*p* across the membrane.

A characterisation of equilibrium shapes of lipid tubules encapsulating macromolecules of different sizes is by coarse-graining tubular shapes observed in simulations, and comparing the deformed shapes with those obtained from a numerical solutions of the shape equations (with surface tension Σ held fixed) to obtain the pressure difference Δ*p* required to bring about the shape change. This pressure difference must exactly balance the entropic pressure exerted on the membrane walls by the polymer. However, due to convergence and stability issues of the numerical scheme used to solve the shape equations (Eq. (4a) - (4c)) for arbitrary values of pressure difference Δ*p*, and surface tension Σ we have not pursued this line of work in our study. Instead, we outline a variational calculation using a shape ansatz outlined below. Our approach turns the functional minimisation of the nonlinear shape equations to a simple function minimisation involving a small number of variables.

## III. VARIATIONAL FORMULATION OF ENERGY MINIMISATION OF TUBULAR SHAPES

We present a variational solution of the shape equation using two ansatz <u>(a)</u> a spherical bulge as shown in Fig. (7b), and <u>(b)</u> and a ellipsoidal bulge on a cylindrical tube as shown in Fig. (11b). Fig. (11b) resembles the geometry of a fluid drop wetting a fibre [31]. The deformed shape is parameterized by the radius of the spherical bulge *R*, and the length *L* and the diameter *D* of the cylindrical tube. Similarly the ellipsoidal bulge is parameterised by the axial deformation scale *ξ*, the maximal radial deformation 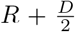, and the length of the tube *L*.

**FIG. 7.**
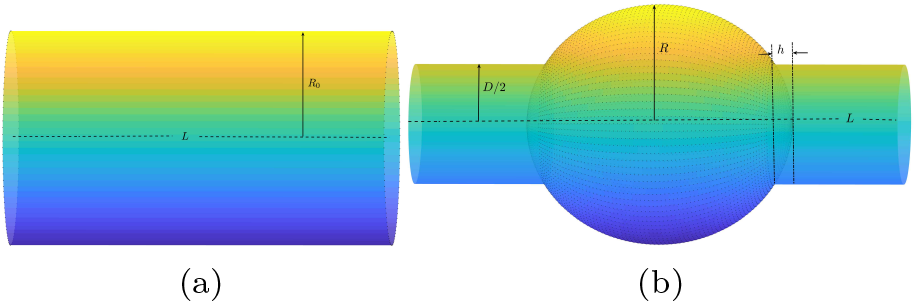
Schematic picture of (a) an undeformed cylindrical tube of length *L* and radius *R*_0_, and a (b) deformed cylinder with a spherical bulge in the centre of radius *R*, and a cylindrical section of diameter *D*. The overlapping region between the cylinder and the spherical bulge is a spherical cap of radius *D*/2 and height *h*.

## A. Spherical bulge model

Fig. (7) shows a deformed tubule with a spherical bulge of radius *R* at its centre on account of the encapsulated polymer. The cylindrical section of the tube has a radius *D/*2 < *R*_0_, such that the total area of the tubule remains the same as that of the undeformed cylinder. The constant area constraint leads to a relation between geometric parameters characterising the deformed and undeformed tubule. Using the parameterisation in Fig. (7b), the total area of the deformed tube with spherical bulge is given by

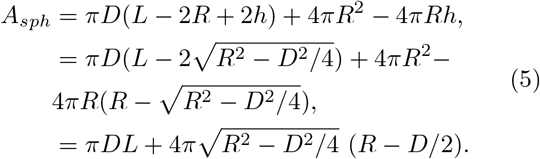

Next, we compute the energetic cost of deforming a cylindrical tubule to form a spherical bulge. Elastic energy of the tubule in the absence of the tensile force and the area and volume constraints is given by,

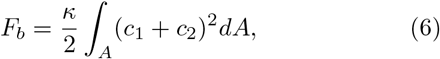

where *c*_1_ and *c*_2_ are the principal curvatures and *κ* is the bending modulus. For sphere of radius *R*, principal curvatures are 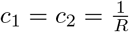, while for a cylinder of radius *D*/2, the principal curvatures are 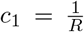 and *c*_2_ = 0 respectively. The bending energy of the deformed tube is given by

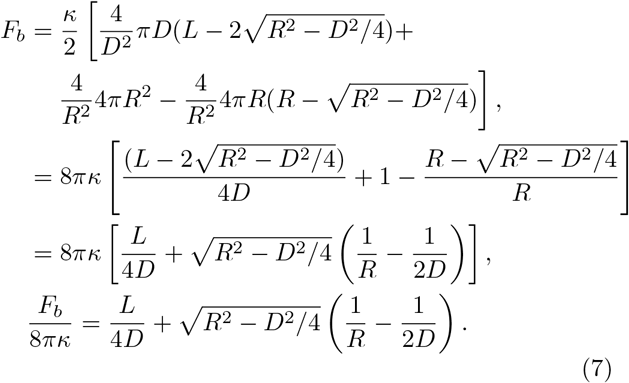

The elastic energy of the undeformed cylindrical tube computed from the Helfrich-Canham free energy Eq. (3) is given by

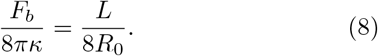

The elastic free energy of the deformed cylinder Eq. (7) approaches that of the undeformed cylinder in the limit, *D*/2 → *R*_0_ and *R* → *R*_0_. This can be seen in Fig. (8a). Further the elastic energy of the deformed cylindrical tubule is always greater than the undeformed one as shown in in Fig. (8a). In equilibrium, the pressure difference Δ*p* across the tubule is zero. Insertion of a polymer chain inside a tubule leads to an excess entropic pressure that changes the volume of the tubule. For a cylindrical tubule with a spherical bulge in the centre to be energetically stable, the sum of the work done in deforming the tubule Δ*pdV* and the elastic energy of the deformed tubule must be equal to the energy of the undeformed cylinder. We numerically obtain the shape parameters *D/R*_0_ and *R/R*_0_ for which the above condition is satisfied.

The work done by the excess pressure Δ*p* on the tubule is given by,

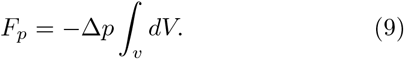

**FIG. 8.**
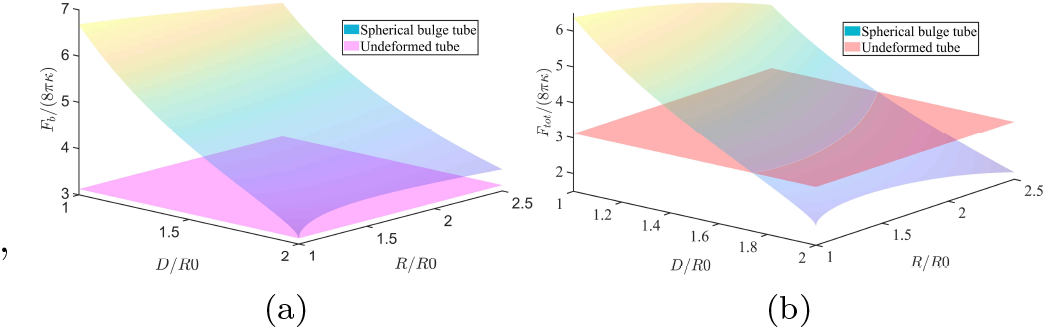
Surface plots of the elastic energy of (a) an undeformed cylindrical tube, compared against a deformed tube with a spherical bulge at its centre. The deformed tubule has a higher bending energy compared to the undeformed cylinder, with the energies being equal when *D/R*_0_ = 2 and *R/R*_0_ = 1. Panel (b) shows the total energy of the deformed cylinder plus the work done by the internal pressure due to chain insertion in increasing the volume of the tubule for 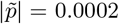, compared against the energy of an undeformed cylindrical tubule. The intersection of the iso-surfaces indicates shape parameters *D/R*_0_ and *R/R*_0_ for the case where the energy of the deformed tube is equal to that of the undeformed tube. The locus of intersection points traces a family of curves that minimise the elastic energy of the tubule subject to the incorporation of the work done by the internal pressure.

For the spherical bulge geometry shown in the Fig. (8a) this yields,

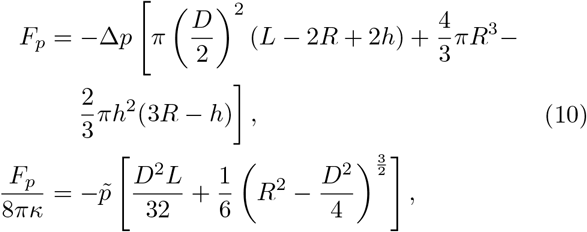

where 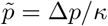. Thus total energy of the deformed tube is sum of bending energy and the work done by the excess pressure, *F*_*tot*_ = *F*_*b*_ + *F*_*p*_. We plot *F*_*tot*_ as a function of the dimensionless parameters *D/R*_0_, and *R/R*_0_. As seen in Fig. (8b) *F*_*tot*_ intersects with lowest energy state of the undeformed tube for a range of pressures 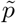. Fig. (8b) shows such an intersection point for 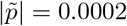.

The intersection between the two planes, i.e., the elastic energy of the undeformed and the deformed tubule traces a curve yielding a family of shapes that minimise the Helfrich-Canham free energy. However, not all solutions satisfy the area constraint. The area of a deformed tubule is plotted in the *D/R*_0_-*R/R*_0_ plane as shown in Eq. (5). The intersection between the two planes i.e., the area of the deformed and undeformed cylindrical tubule traces out yet another family of curves satisfying the constant area constraint as shown in Fig. (9). Implementing constant area constraint in conjunction with the minimisation of the elastic energy we obtain a deformed tubular profile for an imposed pressure 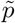. The shape parameters *D/R*_0_ and *R/R*_0_ are uniquely determined for a given pressure 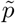.

**FIG. 9.**
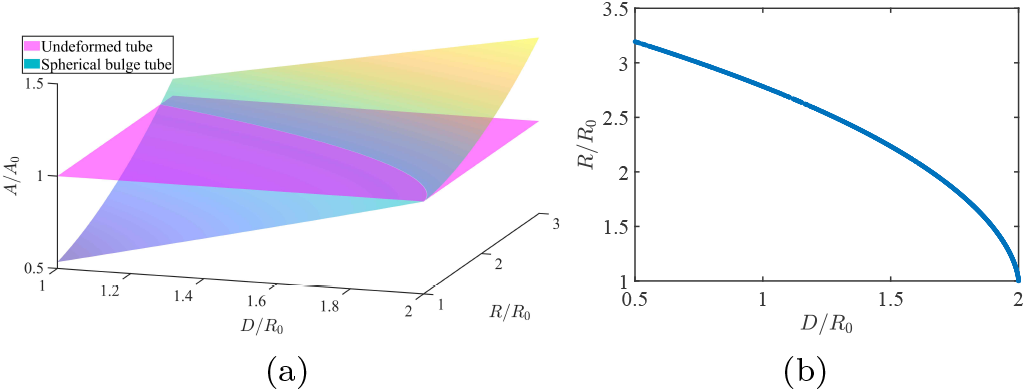
(a) Surface plots of the area of spherical bulge and undeformed cylindrical tube in the *D/R*_0_-*R/R*_0_ plane. The iso-surfaces intersect at points satisfying the constant area constraint of the tubule. The locus of the intersection points of the two iso-surfaces traces a family of curves that satisfy the constraint for different shape parameters as shown in panel (b).

Fig. (10a) shows the intersection point of three planes formed by the elastic energies of the deformed and undeformed cylinder and the area as a function of the variational parameters *D/R*_0_, and *R/R*_0_. The intersection of all three planes yields the unique value of shape parameters for a given pressure 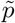. Fig. (10b), shows 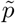 as a function of *R/R*_0_, the radius of the spherical bulge and the diameter of the tube *D/R*_0_ normalised by the radius of the undeformed tubule. As shown in the figure, the equilibrium shape parameters *D/R*_0_ = 2, and *D/R*_0_ = 1 in the absence of pressure correspond to the undeformed cylinder (Fig. (**??**)). The geometric parameters *R/R*_0_ and *D/R*_0_ values can be obtained for a particular choice 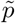, that minimises the elastic free energy obeying the constant area constraint of the deformed membrane tube.

**FIG. 10.**
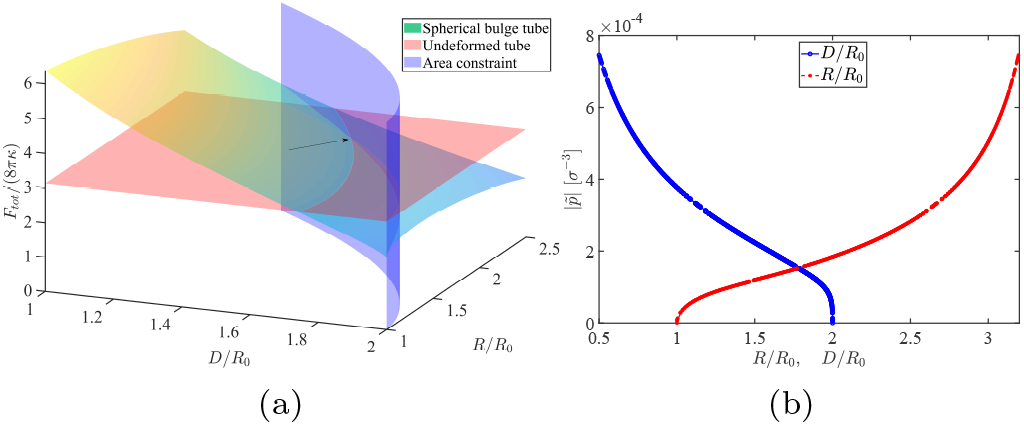
(a) Surface plot of total energy of the deformed cylinder acounting for the work done by the increase in internal pressure due to the insertion of the polymer, for 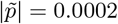 compared against the energy of an undeformed cylindrical tubule. Area constraint will give perticular value of *D/R*_0_ and *R/R*_0_ for that particular 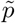. (b) To get minimum energy of the deformed tubule, each 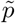 corresponds to particular solution of *D/R*_0_ and *R/R*_0_.

## B. <u>Drop on a fibre</u> or sinusoidal bulge model

Though the variational calculation using a “cylindrical tubule with a spherical bulge” ansatz gives a measure of the excess pressure exerted by the polymer it has a limitation. The slope discontinuity of the tubular membrane at the intersection of the spherical bulge and the undeformed cylinder is unrealistic. We consider a shape ansatz akin to a “drop wetting a fibre” geometry where the deformed section smoothly connects to the undeformed tubule [31] given by,

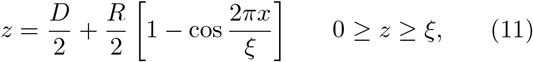

where *z*(*x*) is the displacement of the membrane from its symmetry axis. The deformed axisymmetric shape smoothly connects with the undeformed segment, <u>i.e.</u> a cylinder of length *L* – *ξ* having a diameter *D*. In order to ensure the continuity of the membrane, we match the value of the function *z*(*x*) and the derivative *z′*(*x*) at the intersection point between the deformed and undeformed segments. Further, we ensure that *z′*(*x* = *ξ*) = 0.

We compute the area of the deformed tube using the functional form of Eq. (11), where 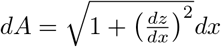

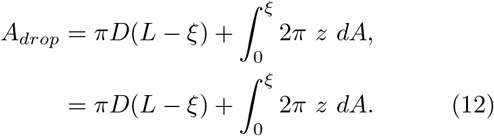

Similarly the volume of the deformed tube using the functional form in Eq. (11) is given by,

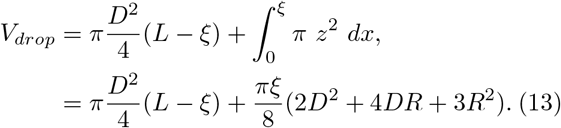

Using Fig. (11b), principal curvatures of the meridian curve and the one perpendicular to the symmetry axis is given by, 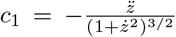, and 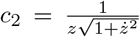 (where 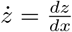 and 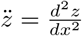) respectively.

**FIG. 11.**
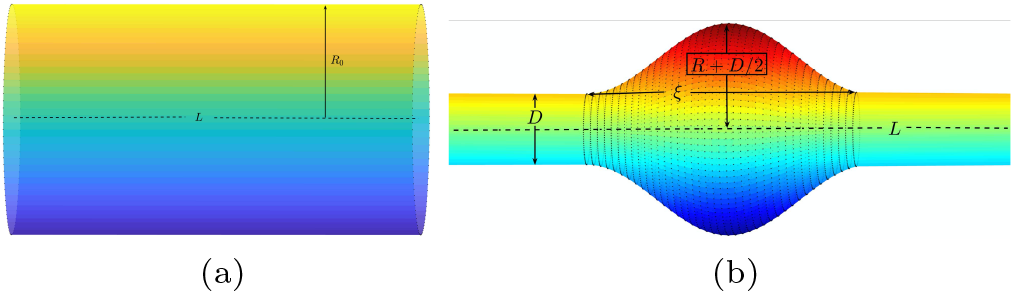
Schematic figure of a tubular membrane showing (a) an undeformed tube, of length *L*, and radius *R*0, and (b) a deformed cylindrical tubule with diameter *D* and length *L*, and a central bulge resembling a “drop wetting a fibre” geometry. We assume an axisymmetric profile. The axial deformation scale of the bulge is *ξ* while the maximum deformation of the tubule is 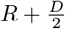.

The elastic energy due to the deformation of the membrane can be calculated from the Helfrich-Canham Hamiltonian [25] and is given by,

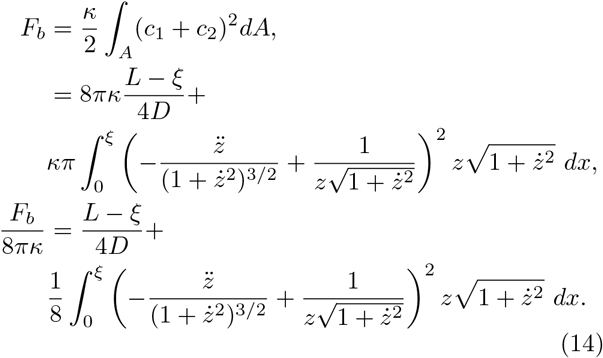

The work done by the pressure in deforming the cylindrical tubule to bring about a shape change is given by Eq. (9)

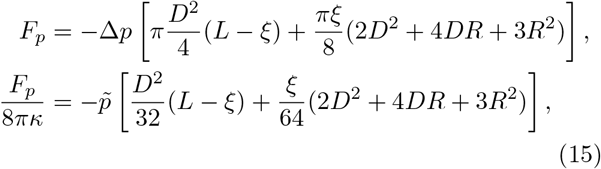

where 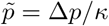.

Fig. (12b) shows the isosurface of the elastic energy of the deformed tubule using the drop on a fibre shape ansatz, and the undeformed cylindrical tubule plotted in the 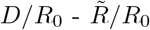 parametric plane 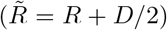, for a fixed value of *ξ*. As in the case for the spherical bulge model the locus of the intersection points between the two iso-surfaces traces out parameter values for which the deformed cylinder has the same energy as an undeformed tubule. Fig. (12b) shows a family of curves for different values of *ξ* and a given value of shape parameters 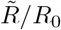, and *D/R*_0_, that minimise the elastic free energy.

**FIG. 12.**
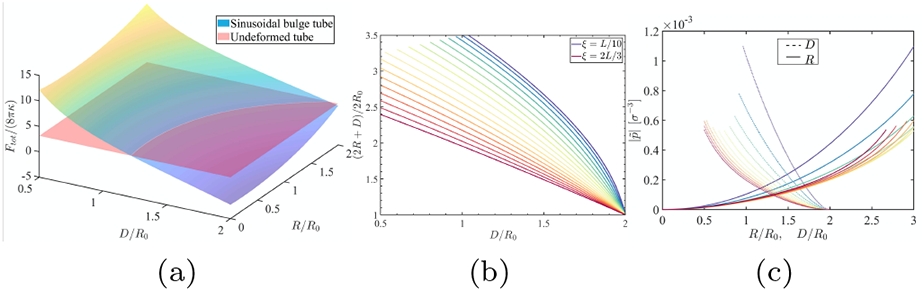
(a) Comparison of the elastic energy of the deformed tubule of length *L* = 300 and *R*_0_ = 12 accounting for the work done by the internal pressure 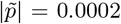, and an undeformed cylindrical tube. The iso-surfaces intersect at a point where the total energies of the deformed and the undeformed tubules are equal. Panel (b) shows a family of curves for different values of *ξ* satisfying the equal energy criterion. The corresponding pressure values 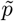, as a function of *D/R*_0_ and *R/R*_0_ for a given value of *ξ* is used to obtain the minimum energy of the deformed tube as shown in panel (c).

In conjunction with the constant area constraint we obtain a unique value of shape parameters *D/R*_0_ and *R/R*_0_ for which the elastic energies of the deformed and undeformed surfaces are equal for a given value of *ξ*. The family of curves for which the area constraint is satisfied is shown in Fig. (12b). The intersection of the family of curves that satisfy equal energy and area constraint between deformed and undeformed tubules gives rise to different pressures 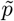.

We note that the three parameter variational minimisation of the elastic free energy is non-unique, as the number of variables *ξ*, 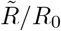, and *D/R*_0_ is more than the number of constraints. It is thus possible to find more than one solution with a different combination of *D/R*_0_, 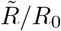, and *ξ* that simultaneously minimise the free energy and satisfy the constant area constraint. We therefore need an additional input to uniquely determine the shape and hence the pressure exerted by an encapsulated polymer chain. This can be carried out in imaging studies by measuring the axial deformation length *ξ*. One can then use the “drop on a fibre” shape ansatz and the calculational scheme outlined here to uniquely determine the pressure exerted by a semiflexible chain on the walls of a soft tubule. Along with the calculation of pressure based on free energy of the confined polymer this forms a self-consistent scheme to connect the conformational states of confined chains with their thermodynamic behavior.

## C. Pressure calculation from polymer statistics

Entropic free energy can be written as, *F* (*N*) = − *k*_*B*_*T* ln *P* (*R*), where 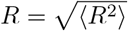, mean end-to-end distance of the confined polymer chain. For confined semi-flexible chains with moderate values of persistence length *l*_*p*_ the chain assumes an ellipsoidal configuration. The volume of the ellipsoid can be obtained a function of chain length *N*, i.e., *V* (*N*). This can be used to compute the pressure exerted by the chain 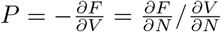. Due to limitations in our system size *N* and long simulation run times required to reach equilibrium we have relegated this to a future study.

## IV. COARSE GRAINED SIMULATIONS

## Description of the Model

To probe the conformational properties of a semiflexible polymer confined in a soft tube, we perform Langevin dynamics simulation to verify theoretical results obtained from continuum theories. For mesoscopic simulations of the tubular membrane, we adopt the coarse-grained <u>Cooke-Kremer-Deserno</u> lipid bilayer model [40]. A coarse grained three bead model of lipid molecules is modelled by a hydrophilic head bead, and two hydrophobic tail beads. In addition to a bonded interaction, head and tail beads interact via a repulsive Weeks-Chandler-Anderson (WCA) or truncated Lennard-Jones(LJ) potential

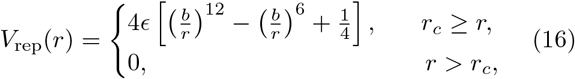

where *ϵ* is the potential well depth and 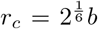 is the location of the minima. The geometric parameters of the three bead lipid model with *b*_head,head_ = *b*_head,tail_ = 0.95*σ* and *b*_tail,tail_ = *σ* is such that the tail beads are slightly bigger than the head bead. To mimic fluidic behaviour of the membrane non-bonded interactions between coarse-grained (CG) tail beads is incorporated. Thus, the effective hydrophobic interaction between tail beads mimicking the implicit solvent is given by

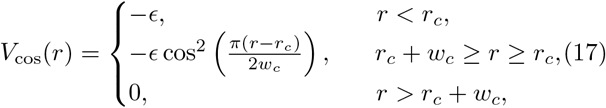

i.e., an attractive potential of depth *ϵ* with the potential decaying to zero smoothly in the interval between *r*_*C*_ and *r*_*c*_ + *w*_*c*_. The width of the attractive region is *w*_*c*_. The bonded interaction between tail beads and head bead is modelled by the finite extensible nonlinear elastic (FENE) potential,

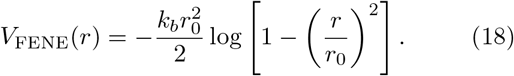

Here *r*_0_ is the maximum extension of the spring.

A bond bending potential, that generates a straight conformation of lipid molecules is given by

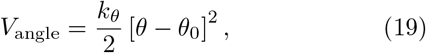

where equilibrium angle *θ*_0_ between three beads is *θ*_0_ = 180° and *k*_*θ*_ is the spring constant corresponding to the bond-bending potential. For lipid interactions *k*_*θ*_ = 10*ϵ*/*σ*^2^, while for the polymer *k*_*θ*_ ranges between 10*ϵ/σ*^2^ and 100*ϵ/σ*^2^ depending on the semiflexibility of the chain.

A <u>Kremer-Grest</u> bead-spring model is used to simulate CG polymer of *N* monomers connected by FENE springs as shown in Eq. (18). The persistence length of the polymer depends upon the equilibrium bond angle *θ*_0_ between three consecutive beads and the spring constant of the angle potential *k*_*θ*_ as outlined in Eq. (19). The beads have diameter *σ* with a bond cut off length *r*_0_ = 1.5*σ*. The monomers interact via the repulsive WCA potential (Eq. (16)). The interaction between monomers and the lipid molecules of the bilayer follow a similar form.

We performed molecular dynamics simulations of polymer-membrane system using the LAMMPS [42] package at a finite temperature using a Langevin thermostat. A semiflexible polymer is inserted inside a cylindrical bilayer membrane. The initial polymer conformations were generated using a MATLAB script. Though self-assembly of a flat membrane has been observed in simulations, spontaneous assembly of a bilayer to form tubules have not yet been observed [40]. We generated a cylindrical bilayer tubule as our initial configuration. We minimise the energy of the compound polymer-membrane system via a conjugate gradient minimisation scheme. To obtain equilibrium conformations for both polymer and membrane we performed simulations in a constant volume ensemble, for *t* = 5 × 10^6^ Δ*t*, where Δ*t* is the time step of the simulation. For all the results reported in the paper, Δ*t* = 10^−2^*τ*_*LJ*_, where 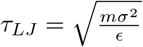, where *m* is the mass of the beads, *σ* is the short distance cut off and *E* is the depth of the Lenard-Jones potential. The coupled polymer tubule system equilibrates after *t*_*eq*_ ≈ 2.5 × 10^6^Δ*t* starting from its initial configuration.

The radius of gyration *R*_*G*_ and the radial *R*_⊥_ and axial extension *R*_∥_ of the polymer is monitored as a function of time. For chains undergoing a conformational collapse the *R*_*G*_ decreases as a function of time. Equilibration time *τ*_*eq*_ is obtained by noting the time scale at which *R*_*G*_ fluctuates around a steady value. The total energy of the system of the system in this phase fluctuates around an average value. For polymers that remain in the swollen phase, we collect data for *t* >> *τ*_*eq*_ obtained from the collapse runs.

## V. DATA ANALYSIS

## A. Quantifying conformational transitions

## <u>Coil to globule</u> transition

A semiflexible polymer inside a bilayer tubule with bending modulus *κ* >> *k*_*B*_*T*, is rigid. As a result the encapsulated polymer is stretched out and adopts a linear conformation. On the contrary, if *κ* ~ *k*_*B*_*T*, polymer can deform the tube and adopt a globular structure. The confined polymer and the bounding tubule adopt different shapes depending on the chain flexibility, its size *N*, bending modulus of the tubule *κ*, and membrane tension Σ. For stiff chains with, *l*_*p*_ >> *R*_0_, the chain adopts a toroidal shape, in order to minimize its bending energy. The resulting tubular shapes are qualitatively different than those obtained for softer chains, <u>i.e.</u> *l*_*p*_ ~ *D*. In this case, *l*_*p*_ ~ *D*, polymer adopts a globular ellipsoidal conformation with the major axis along the tube axis. The axisymmetric shapes are quantified by computing the eigenvalues of gyration tensor with the maximum eigenvalue being along the tube axis. The eigenvalues along the two orthogonal principal axis in the radial plane have similar values. Prolate ellipsoidal shapes obtained in this regime are shown in the main text Fig. (4).

Fig. (13) shows the variation of the radius of gyration *R*_*G*_(*t*) of a semiflexible polymer chain in an extended initial conformation placed inside a bilayer tubule undergoing a coil to globule transition. The initial radius of gyration of the polymer chain of size *N* = 2000 is *R*_*G*_(0) ≈ 58*σ*. The time trace shows *R*_*G*_(*t*) decreasing as a function of time indicating the collapse transition of the chain. Configuration snapshots indicate a transition to a final ellipsoidal globular structure. The polymer-tubule system is simulated for *τ* ≈ 10^7^Δ*t*, with data collected after *τ* ≈ 5 × 10^6^Δ*t*, when the system has equilibrated. The data is collected in intervals of *τ*_*m*_ = 10^3^Δ*t*.

**FIG. 13.**
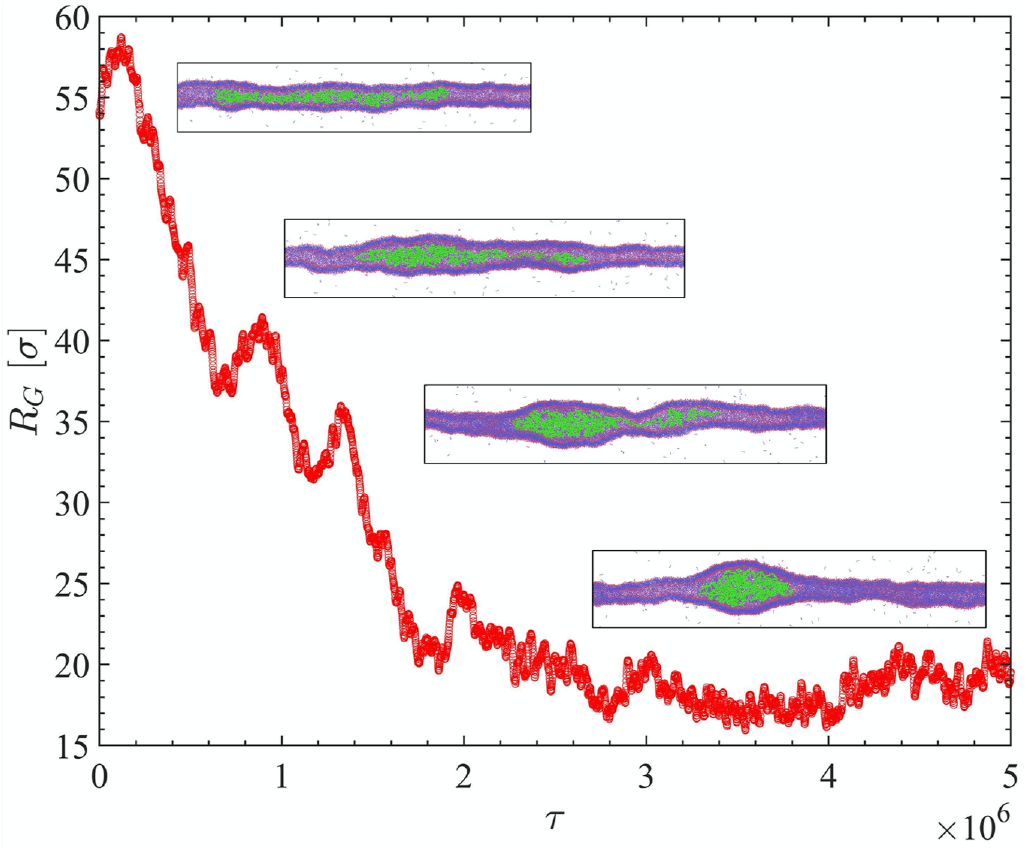
Variation of the radius of gyration of the polymer chain *RG* as a function of time undergoing a collapse transition from an extended <u>coil</u> to <u>globular</u> state. Conformational statistics data is collected after *τ* ≈ 5 × 10^6^Δ*t* when the chain equilibrates every *τ*_*m*_ = 10^3^Δ*t* time steps. Configuration snapshots reveal the dynamics of chain collapse including presence of metastable states.

As expected the <u>coil</u> to <u>globule</u> transition of the chain also depend on the mechanical properties of the confining tubule. In the limit of a rigid tube *κ* >> *k*_*B*_*T*, polymer prefers to be in an extended state. The bending modulus of the membrane can be tuned in simulations by changing the strength of the attractive interaction between the tail beads. This can be effected by either changing the range of the potential *w*_*c*_/*σ* and the effective temperature *k*_*B*_*T/ϵ*. We have obtained a phase diagram to explore <u>coil</u> to <u>globule</u> transition in the *w*_*c*_/*σ* −*kT/ϵ* plane. The final configuration snapshots after *t* = 5 × 10^6^ time steps are shown in the Fig. (14). Corresponding temperatures *k*_*B*_*T*, bending modulus of the tubule, *κ* and the averaged tube radius squared 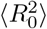 are shown in the Table (I). The bilayer tubule is not stable at temperatures higher than those shown in the table (I).

**TABLE I.** Bending rigidity *κ*, and square averaged tubular radius 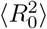 for tubules shown in Fig. (14) without an encapsulated polymer. The number of lipids used in simulating the tubules are *N*_*t*_ = 97200.

| Tube properties | $w_c = 1.3\sigma$ | $w_c = 1.4\sigma$ | $w_c = 1.5\sigma$ |
| --- | --- | --- | --- |
| (i) $k_B T/\epsilon$<br>$\kappa/k_B T$<br>$\langle R_0^2 \rangle/\sigma^2$ | | | 0.9<br>28.7667<br>79.8670 |
| (ii) $k_B T/\epsilon$<br>$\kappa/k_B T$<br>$\langle R_0^2 \rangle/\sigma^2$ | | 0.9<br>15.7465<br>104.3290 | 1.0<br>18.4059<br>102.4582 |
| (iii) $k_B T/\epsilon$<br>$\kappa/k_B T$<br>$\langle R_0^2 \rangle/\sigma^2$ | 0.8<br>14.2495<br>106.9920 | 1.0<br>14.1763<br>119.2689 | 1.1<br>16.4098<br>114.9923 |
| (iv) $k_B T/\epsilon$<br>$\kappa/k_B T$<br>$\langle R_0^2 \rangle/\sigma^2$ | 0.9<br>11.9555<br>124.1086 | 1.1<br>11.3364<br>133.5598 | 1.2<br>12.3305<br>126.8585 |
| (v) $k_B T/\epsilon$<br>$\kappa/k_B T$<br>$\langle R_0^2 \rangle/\sigma^2$ | 1.0<br>9.9009<br>141.1741 | 1.2<br>5.6646<br>144.8585 | 1.3<br>7.9215<br>135.7489 |

**FIG. 14.**
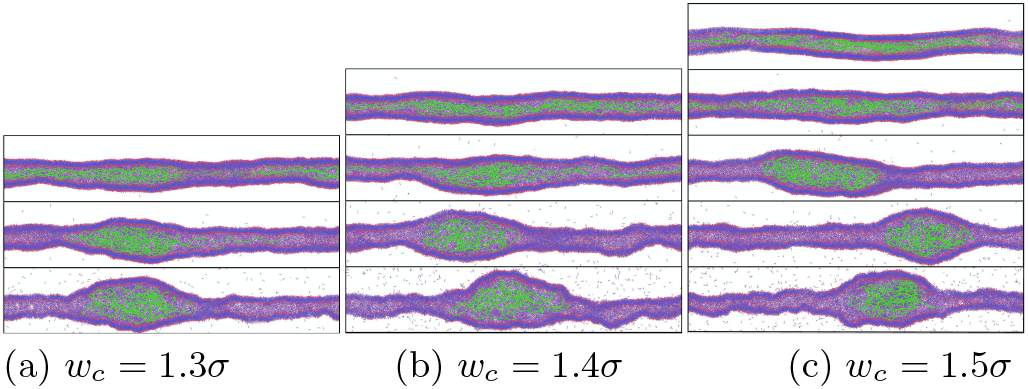
Simulation snapshots of extended coil and globular equilibrium polymer conformations inside a bilayer tubule with different bending moduli *κ* and surface tension Σ obtained by varying the range of the lipid interaction potential *wc* among tail beads and temperature *k*_*BT/ϵ*_. The data is collected after *t* = 5 × 10^6^Δ*t* when thermal equilibrium is established.

## B. Statistical properties of Tubule

The bilayer tubule used in our simulations has a finite thickness, with the outer and inner diameters being *D*_*out*_ and *D*_*in*_ respectively. In order to determine the radial bulge of the tubule we divide the cylindrical tubule in slices of width *σ*, <u>i.e.</u>, the size of a head bead. Therefore the tubule can be visualised as a series of adjacent rings of width *σ*. The instantaneous tube diameter at position *x* is calculated by taking the mean of the inner and outer *D*(*x*) = (*D*_*in*_(*x*) + *D*_*out*_(*x*))/2 diameters. For a fluctuating tubule *D*_*out*_ and *D*_*in*_ are the distance of the head beads from the centre of mass of each ring. We compute ⟨*D*(*x*)⟩ by performing a time average of the instantaneous tube diameter measured every *τ*_*m*_ = 10^3^Δ*t* time steps after equilibration *τ*_*eq*_. For axisymmetric tubular shapes ⟨*D*_*in*_(*x*)/2⟩ and ⟨*D*_*out*_(*x*)/2⟩ is independent of the azimuthal angle. This is seen for the tubule without a confined polymer and for ellipsoidal tubular shapes. However for toroidal coils *l*_*p*_ >> *R*_0_ the azimuthal symmetry is lost. In this case we average over the azimuthal angle for all head beads in a given ring along the tubule, to obtain instantaneous values of the tubule diameter *D*_(_*x*) and in turn its thermal average ⟨*D*(*x*)⟩. Fig. (15a) shows the time averaged radial bulge of an axisymmetric tubule along the tube axis having a confined polymer. The outer and inner inner leaflets ⟨*D*_*out*_/2⟩ and ⟨*D*_*in*_/2⟩ are shown by dashed lines. The area of the tubule remains constant throughout the simulation (Fig. (15b)) while conformational transition of the chain is accompanied by a volume increase (Fig. (15c)).

**FIG. 15.**
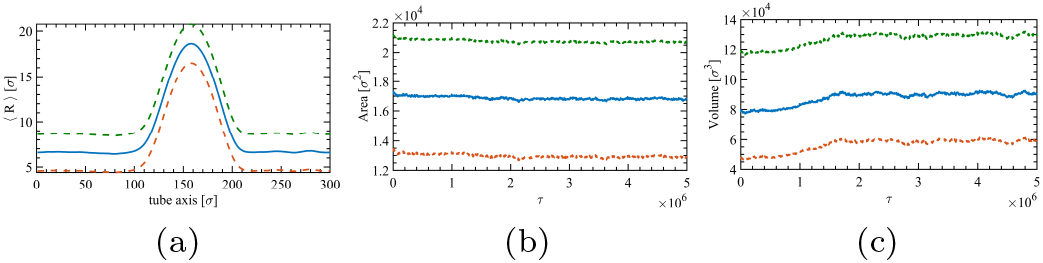
<u>(a)</u> Radial bulge along the axis of the tube *x* for outer ⟨*D*_*out*_/2⟩ (dashed orange), inner ⟨*D*_*in*_/2⟩ leaflets (dashed purple), and the <u>i.e.</u> average radius ⟨*D*(*x*)/2⟩. Panel <u>(b)</u> shows that the area of the bilayer tubule remains constant over time. Panel <u>(c)</u> shows that the volume of the tube increases during the coil to globule transition.

The centre of mass of each ring of lipid molecules that form the tubule is allowed to fluctuate. Fig. (16)(right panel) shows the radial deformation profile along the tube axis as a function of time. Due the motion of the centre of mass of the deformed segment of the tubule along the tube axis (SI movies) the maximum radial bulge occurs at different positions *x*. A simple averaging of the deformed configurations over time to compute the maximum value of the radial bulge, ⟨*D*_*m*_/2⟩ would therefore be erroneous. Therefore in order to compute the radial bulge (and since we have periodic boundary conditions imposed along the direction of the tube) we shift the deformed tubular profile, along the tube axis such that the maximum deformation occurs at *x* = *L*/2. Fig. (17)(a) shows the variation of the maximum radial bulge of the tubule as a function of the polymer size *N*. The maximum radial bulge increases with *N* as ⟨*D*_*m*_/2⟩ ~ *N*^0.27^ (see Fig. 2 of the main text). In order to obtain the spatial deformation scale along the tube axis we compute the numerical derivative of the radial deformation profiles, <u>i.e.</u>, 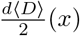 (Fig. (17)(a)). The extremities of the zero crossing of this curve determines the axial deformation scale *ξ*. As shown in Fig. (17)(b) *ξ* is independent with polymer size *N*. We have observed similar behaviour for all values of initial tube radius *R*_0_.

**FIG. 16.**
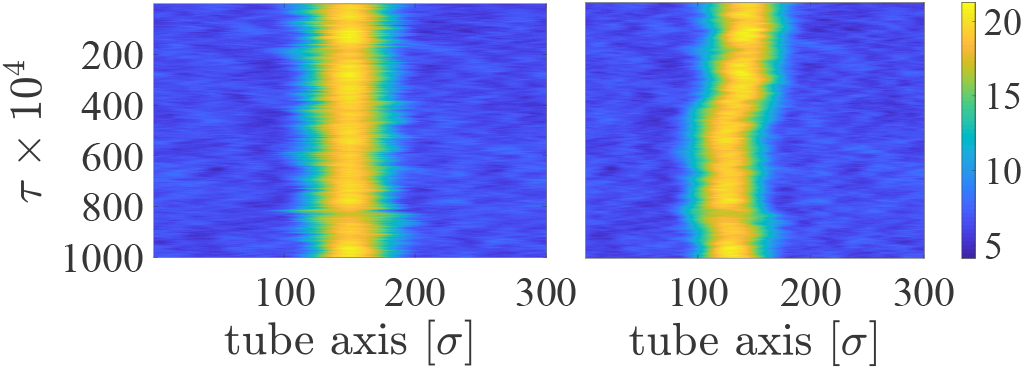
Space time plots of the radial bulge of a deformed tube along the tube axis (right). Colormap indicates the value of the radius of each tubular cross section (ring of lipid molecules). In order to compute the time average the deformed profile at each time instant is shifted along the tube axis such that the maximum deformation (brightest point) occurs at *x* = *L*/2 (left).

**FIG. 17.**
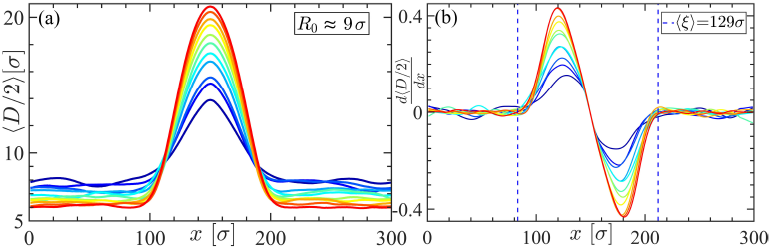
Radial bulge ⟨*D*(*x*)/2⟩ of a bilayer tubule of initial radius *R*_0_ ≈ 9*σ*, along its axis *x*, encapsulating a semiflexible polymer as a function of its size *N*. The polymer size is varied between *N* = *N*_*min*_ = 600 (blue line) to *N*_*max*_ = 2600 (red line) in increments of Δ*N* = 200 (a) obtained from coarse-grained simulations. The maximum radial bulge ⟨*D*_*m*_/2⟩ in-creases with *N* (panel (a)), while the axial deformation length *ξ* is independent of the chain size (panel (b)). The radius of undeformed cylindrical segments is less than *R*_0_ to maintain the constant area constraint.

## C. Statistical properties of Confined Chains: long length scales *l* ~ *ξ*

The statistical properties of the polymer chain in the long wavelength limit *l* ~ *ξ* are calculated by noting the position of the terminal monomers of the deformed chain. These include, ⟨*R*_⊥_⟩, the maximum extent of the polymer along radial direction and ⟨*R*_∥_⟩ the maximum extent of the polymer along the tube axis. In order to compute ⟨*R*_⊥_⟩ we average over the radial coordinates <u>i.e.</u>, the *xy* plane. Monomer position snapshots are collected every *τ_m_* = 10^3^Δ*t* time steps once equilibrium *t* = *τ*_*eq*_ is reached.

## D. Statistical Properties of Confined Chains: short distance *l*_*p*_ ~ *l* ≲ *ξ*

## 1. *Asphericity* Δ *and the nature of asphericity (*Ξ)

Confined polymer shapes over short length scales are quantified by two parameters, the asphericity (Δ) and the nature of asphericity (Ξ) [34].These quantities are computed from the gyration tensor (see main text). Intuitively, the parameter Δ is related to the variance of the three eigenvalues 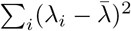, where *λ*_*i*_ = *λ*_*x*_, *λ*_*y*_, *λ*_*z*_ and 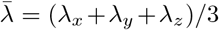, while nature of asphericity Ξ is related to the product of the eigenvalues 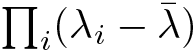. To visually describe the shapes, we rescale these two parameters 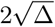 ranges [0 2] and cos^−1^ Ξ/3 ranges [0 *π*/3] [34]. The geometric interpretation of the shapes is shown in Fig. (4) (in the main text). The value Δ = 1, and Δ = 0 corresponds to a rod and sphere respectively, while the parameter, Ξ is used to characterise prolate vs. oblate shapes.

## 2. *Pair correlation function g*(*r*)

The pair correlation function or radial distribution function *g*(*r*) describes the variation of the density of monomers as a function of distance from a reference particle. Thus *g*(*r*) is the ratio between the average number density at a distance *r* from any given monomer as shown in the Fig. (18). The pair correlation function for spherical and tubular confinements of polymer chains with different stiffness (corresponding to Fig. (4) of the main text) are shown in Fig. (18). The position of the maxima of the radial distribution function *g*(*r*) corresponds to the length scale associated with the order in the system. As shown in Fig. (18) both the peak positions as well as the behaviour of *g*(*r*) for large *r* is dependent on the nature of the confinement and the mechanical properties of the chain. Thus *g*(*r*) can be used to characterised confined polymer conformations and can be probed via scattering experiments by measuring the static structure factor *S*(*q*). A full exploration of the static and dynamic structure factors, *S*(*q*), and *S*(*q, ω*) is relegated to a future study [43].

**FIG. 18.**
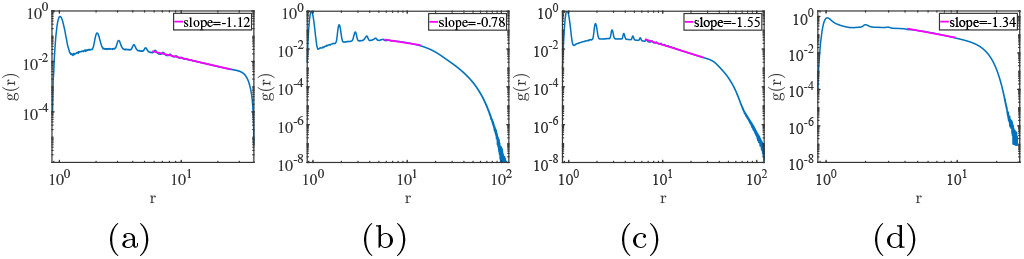
Pair correlation function for compact polymer conformations inside a soft tubule corresponding to different confining geometries, and chain stiffness <u>e.g.</u>, (a) inside a rigid sphere *l*_*p*_ ≈ 200*σ*, (b) soft tubule with *l*_*p*_ ≈ 13*σ*, (c) *l*_*p*_ ≈ 200*σ*, and (d) *l*_*p*_ ≈ 200*σ* in a bad solvent (as in Fig. (4) of the main text). The positions of the peaks in *g*(*r*) and its decay at large distances depends on the chain stiffness and the confining geometry and can be used to characterise compact confined polymer conformations.

## 3. *Radial monomer density ρ*(*r*)

Density distribution of a polymer chain in different environments averaged over *τ*_*m*_ = 5000Δ*t* time steps is shown in Fig (4) of the main text. We compute the distance of a monomer *r* from the centre of mass coordinate 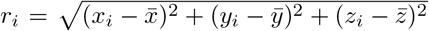. A histogram is obtained by binning the monomers as a function of *r* which gives the density distribution *ρ*(*r*). In case of strong confinement, the monomer density is high near the boundary, while for weak confinement the monomer distribution is peaked about the centre of the channel. The monomer density is used to distinguish between ellipsoidal and toroidal coils as stated in the main text.

## VI. SI: MOVIE

SI movie shows the time evolution of a polymer encapsulated in a tubule starting from a random initial polymer conformation to the equilibrium sate obtained using CGMD simulations. The first and second columns show the axial and radial cross sections of lipid tubule. The encapsulated semiflexible polymer (green beads) of size *N* = 2000*σ*, is confined in the bilayer lipid tubule having hydrophilic head groups (red beads) and hydrophobic tails (blue beads). For rigid tubules having *κ* = 24*k*_*B*_*T* (a) swollen chain is seen (first row), while for softer *κ* = 12*k*_*B*_*T* tubules (second row) (b) a prolate ellipsoidal conformation is observed. The persistence length of the polymer in this case is *l*_*p*_ = 13*σ*. The third row (c) shows a toroidal coil for a polymer with *l*_*p*_ ≃ 200*σ*, (such that *l*_*p*_ ≳ *ξ*).

